# Fetal brain response to maternal inflammation requires microglia

**DOI:** 10.1101/2024.02.21.581282

**Authors:** Bridget Elaine LaMonica Ostrem, Nuria Dominguez Iturza, Jeffrey Stogsdill, Tyler Faits, Kwanho Kim, Joshua Z. Levin, Paola Arlotta

**Affiliations:** Department of Stem Cell and Regenerative Biology, Harvard University, Cambridge, MA, USA; Department of Neurology, University of California, San Francisco, San Francisco, CA, USA; Stanley Center for Psychiatric Research, Broad Institute of MIT and Harvard, Cambridge, MA, USA; Klarman Cell Observatory, Broad Institute of MIT and Harvard, Cambridge, MA, USA

## Abstract

*In utero* infection and maternal inflammation can adversely impact fetal brain development. Maternal systemic illness, even in the absence of direct fetal central nervous system infection, is associated with an increased risk of autism and schizophrenia in affected offspring. The cell types mediating the response of the fetal brain to maternal inflammation are largely unknown, hindering the development of therapies to prevent and treat adverse neuropsychiatric outcomes. Here, we show that microglia, the resident phagocytes of the brain, are enriched for expression of receptors for relevant pathogens and cytokines, throughout embryonic development. Using a rodent maternal immune activation (MIA) model in which polyinosinic:polycytidylic acid is injected into pregnant dams, we demonstrate long-lasting transcriptional changes in fetal microglia that persist into postnatal life. We find that MIA induces widespread gene expression changes in neuronal and non-neuronal cells; importantly, these responses are abolished by selective genetic deletion of microglia, indicating that microglia are required for the transcriptional response of other cortical cell types to MIA. These findings demonstrate that microglia play a critical, durable role in fetal response to maternal inflammation, pointing at microglia as a potential therapeutic cell target.

## INTRODUCTION

Maternal infections and inflammation can disrupt fetal brain development and lead to significant neurological and psychiatric problems in affected offspring^1,2^. While some pathogens directly infect fetal brain cells, widespread maternal immune system activation and cytokine release in the absence of direct central nervous system (CNS) infection in the fetus are also detrimental to neurodevelopment^3^. Maternal fever, elevated serum cytokine levels, and infection with pathogens that do not cross the placenta, increase the risk of problems such as schizophrenia and autism in affected offspring^1,4–7^. The molecular mechanisms that underlie this association are not well understood.

Maternal immune activation (MIA) in rodents is a well-characterized model for *in utero* inflammation in which pregnant mice are injected with an immune stimulant such as polyinosinic:polycytidylic acid (polyI:C), a double-stranded RNA molecule that mimics viral infection^8^. The offspring of mice subjected to MIA display autism-like phenotypes that are most pronounced in male mice, including abnormal ultrasonic vocalizations and altered interactions with novel objects and mice^9^. MIA affects dendritic spine density and cortical lamination and leads to global changes in protein expression in the developing cortex via activation of the integrated stress response^10–12^. However, the specific cellular targets of MIA in the developing brain that mediate autism-like behavioral phenotypes observed in juvenile and adult mice are unknown.

Microglia, the resident phagocytic cells of the nervous system, are poised to respond to a variety of early insults, and express receptors for some of the cytokines implicated in MIA, including interleukin-17a (IL-17a) and interleukin-6 (IL-6)^13,14^. Microglia originate from myeloid precursors in the yolk sac and begin to populate the embryonic mouse cortex around embryonic day 9 (E9)^15^. Recent studies have demonstrated that microglia play a significant role in the cellular pathophysiology of congenital brain infection with Zika virus and cytomegalovirus and that MIA impacts microglial motility and the ability to respond to immune stimuli in adulthood^16–21^. However, it is unknown how MIA affects microglial gene expression *in* utero or whether microglia are necessary for the transcriptional changes observed across other embryonic cell types in the developing brain in response to MIA^12^. Here we show that MIA alters embryonic and juvenile microglial gene expression profiles, and that microglia are necessary for mediating the effects of maternal inflammation on neighboring neurons and neuronal progenitors in the fetal cerebral cortex.

## RESULTS

### Pathogen and cytokine receptor expression across cell types in the developing cerebral cortex

We sought to confirm the expression of receptors relevant to maternal infection and inflammation in microglia and to identify other cell types that may directly respond to these stimuli. We first interrogated a previously-generated single-cell RNA sequencing (scRNA-seq) dataset for expression in embryonic cerebral cortex of receptors for pathogens known to cross the placenta and cause fetal infection (termed TORCH pathogens, and of receptors for cytokines involved in the inflammatory response to maternal infections^13,14,22–28^ (Fig. 1A). To generate this dataset, the full thickness of the developing somatosensory cortex was isolated and all embryonic cortical cell types were profiled with high temporal resolution from E12.5 through P1^22^. We found that receptors implicated in MIA were robustly expressed in microglia, and absent in progenitor and neuronal cell types, including *Tlr3*, the receptor for Poly(I:C), and *Il17ra*, the receptor for IL-17a, which plays a critical role in mediating behavioral phenotypes in MIA^13^. We calculated a gene module score to quantify cumulative expression of TORCH and cytokine receptors over time, henceforth referred to as CytoTORCH module score (Fig. 1B and Methods). Microglia displayed a higher score than any other cell type in the developing cortex (p<0.05 for Analysis of Variance [ANOVA] test and for pairwise comparisons between microglia and all other cell types using Tukey’s Honest Significance tests [HST]). Several non-neuronal cell types, including pericytes and endothelial cells, also displayed a relatively high CytoTORCH module score. Overall, these data suggest that microglia are poised to act as immediate responders to a broad range of inflammatory and infectious insults in the developing cortex.

**Fig. 1.**
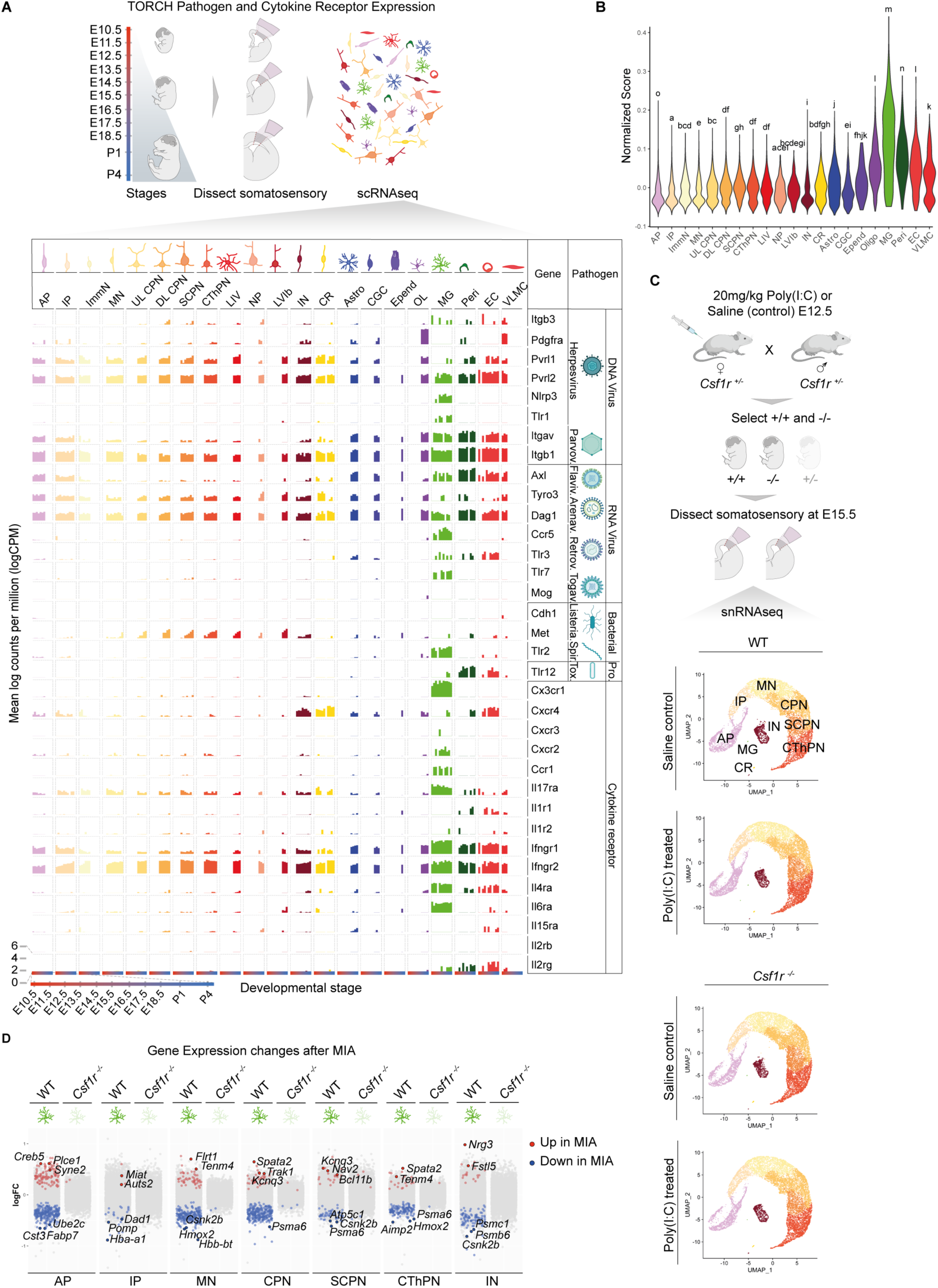
Brain transcriptional changes after MIA require microglia. (A) Simplified schematic of sample processing pipeline from [Di Bella 2021] and bar plot of TORCH pathogen and cytokine receptor expression (mean log counts per million) in each cell type across development. (B) Violin plot of CytoTORCH module score for each cell cluster (MG p<0.05 for ANOVA and for pairwise comparisons between microglia and all other cell types using Tukey’s HST). (C) Overview of experimental design (up) and UMAP visualization of snRNAseq data from the developing somatosensory cortex of E15.5 WT and *Csf1r^-/-^* mice after saline or Poly(I:C) maternal injection at E12.5 (2-3 embryos per condition from a total of 4 pregnant dams, 42,736 total cells) (down). Cells are colored by cell type assignment. (D) Dot plot of log(fold change) in gene expression between MIA and saline for cell types with significant changes at E15.5. DEG analysis performed using NEBULA. AP, apical progenitors; IP, intermediate progenitors; ImmN, immature neurons, MN, migrating neurons; (UL/DL) CPN, (upper layer/deep layer) callosal projection neurons; SCPN, subcerebral projection neurons; CThPN, corticothalamic projection neurons; LIV, layer IV stellate neurons; IN, interneurons; CR, Cajal-Retzius cells; Astro, astrocytes; CGC, cycling glial cells; Epend; ependymal cells; OL, oligodendroglia; MG, microglia; Peri, pericytes; EC, endothelial cells; VLMC, vascular and leptomeningeal cells. Illustrations created with BioRender.com.

### Microglia mediate the response to immune activation of other cell types in the developing cortex

To determine whether microglia directly impact how other embryonic neocortical cell types respond to maternal inflammation, we performed single-nucleus RNA sequencing (snRNA-seq) of the developing somatosensory cortex in *Csf1r* null mice and wild-type (WT) littermate controls after MIA or sham saline injection at E12.5 (Fig. 1C). *Csf1r* null mice have grossly normal embryonic brain development but lack microglia, as the encoded receptor, CSF1R, is required for microglial development^29^. Microglia are the only cell types in the CNS that express CSF1R; therefore, the brain effects of *Csf1r* knockout can be attributed to the loss of microglia. We focused our analysis on the developing somatosensory cortex, due to the previous detailed developmental characterization of the region’s cell types and the demonstrated abnormalities in the somatosensory cortex in the setting of MIA and autism^30,31^. We analyzed male mice only, given the sex-dependent effects of MIA at both the molecular and behavioral level^32,33^. An appropriate maternal inflammatory response to Poly(I:C) injection was confirmed via cytokine array profiling (Fig. S1A). Gene expression was analyzed at E15.5, to allow sufficient time (72 hours) for microglia to express cell signaling factors that may affect surrounding cell types. After quality control and doublet filtering (Table S1 and Methods), we analyzed the expression profiles of 42,736 cells. We clustered the cells using the Louvain algorithm and annotated the clusters *post hoc* based on expression of canonical marker genes (Fig. S1B). We identified 16 clusters, with identities and transcriptional profiles corresponding to 9 previously described cell types at E15.5^22^, including apical progenitors (APs) and intermediate progenitors (IPs), migrating neurons, major subtypes of cortical projection neurons and interneurons, and Cajal Retzius cells (Fig. 1C). As expected, we observed a small cluster representing microglia in WT animals; this cluster was nearly abolished in *Csf1r* null mice (Fig. S1C). The absence of microglia in *Csf1r* null mice was confirmed by immunohistochemistry (Fig. S1D). MIA did not alter non-microglial cell type distributions in WT or *Csf1r* null mice (binomial generalized linear mixed effects model, Fig. S1C).

Differentially expressed genes (DEGs) between MIA and saline control for each cell type were identified by Negative Binomial mixed model Using a Large-sample Approximation, or “NEBULA”^34^. In agreement with previous findings^12^, MIA led to significant gene expression changes across multiple cell types in the developing cortex (Fig. 1D). We found that eliminating microglia via *Csf1r* knockout abolished most of these effects. Similar results were obtained by analyzing DEGs via DESeq2 using pseudobulked counts (Fig. S1E and Methods).

We examined MIA-induced, microglia-dependent gene expression changes in neurons and neuronal progenitors. We found several expression changes in genes involved in neuronal development (Table S2). For example, in WT but not *Csf1r* null mice exposed to MIA, migrating neurons showed increased expression of *Flrt1*, which encodes a cell-cell adhesion molecule that regulates neuronal migration and neurite outgrowth^35,36^. Migrating neurons, corticothalamic projection neurons (CThPNs), and callosal projection neurons (CPNs) showed increased expression of *Tenm4*, a risk gene for schizophrenia that encodes a transmembrane protein that regulates axon guidance^37,38^. APs showed increased expression of *Auts2*, a marker of frontal cortical identity that is involved in synapse formation and axon elongation^39^. We performed Gene Ontology (GO) analysis of DEGs using the enrichGO() function from clusterProfiler to more broadly query molecular pathways and functions that are altered by *in utero* inflammation in a microglia-dependent manner. GO analysis demonstrated that immune activation led to microglia-dependent upregulation of genes involved in neuronal fate commitment, axon guidance, and non-canonical Wnt signaling (Fig. S1F).

We observed that MIA in the presence, but not absence, of microglia induced widespread down-regulation in expression of genes involved in proliferation, energy metabolism, and protein catabolism (Fig.S1G). The expression of multiple proteasomal subunits was reduced, including *Psma6* (migrating neurons, CThPNs, CPNs, APs, interneurons), *Psmb6* (migrating neurons, CThPNs, CPNs, APs, interneurons), *Psmb2* (APs, migrating neurons), and *Psmb4* (CPNs, APs). There was also a reduction in hemoglobin gene expression, including *Hbb-bs* (CPNs, APs, IPs), *Hbb-bt* (APs, IPs, migrating neurons, CPNs), *Hba-a1* (CPNs, APs, IPs, migrating neurons), and *Hba-a2* (APs, migrating neurons), suggesting widespread alterations in cell energetics. Furthermore, MIA reduced the expression of genes involved in positive regulation of viral processes in WT but not *Csf1r* null mice. For example, in CThPNs, CPNs, and migrating neurons, MIA reduced expression of *Tsg101*, which encodes an ESCRT complex member that mediates viral budding, while expression of *Banf1*, which is involved in retroviral intermolecular integration, was reduced in APs and CPNs^40,41^. Additionally, in APs, IPs, SCPNs, CThPNs, CPNs, migrating neurons, and interneurons, MIA resulted in downregulation of *Ppia*, which functions in the replication and infectivity of several viruses and regulates the host type I interferon response to viral infections^42^. All of these responses were abrogated in the *Csf1r* null mice. Taken together, these findings show that microglia are necessary for widespread gene expression changes that occur in multiple embryonic CNS cell types in response to maternal inflammation.

### Single-cell transcriptional atlas of embryonic microglia

To fully delineate microglial molecular identity in the developing somatosensory cortex and to ask whether specific microglial substates could be responsible for mediating the effects of MIA, we built a comprehensive scRNA-seq atlas of purified microglia across embryonic cortical development. Prior studies of microglial transcriptional heterogeneity have included limited embryonic ages and brain regions, precluding a detailed understanding of lineage relationships and substate specific roles in development and disease in the context of dorsal cortical development^43–45^. We performed scRNA-seq on WT microglia isolated from the prospective mouse somatosensory cortex throughout cortical development: embryonic day (E)12.5 and E13.5 (birthdate of layer 5 and 6 excitatory neurons), E14.5, E15.5, and E16.5 (birthdate of layer 2/3 and 4 excitatory neurons), E18.5 and postnatal day (P)1 (end of neurogenesis, transition to gliogenesis). To reduce *ex vivo* microglial activation, we performed nonenzymatic single-cell isolation under cold conditions^43,46^ (Fig. 2A and Methods). Prospective mouse somatosensory cortical tissue was minced and dounce-homogenized, and microglia were labeled with antibodies against CX3CR1, CD11b, and CD45 and purified via fluorescence-activated cell sorting (FACS) (Fig. S2A). Overall, we collected 831 to 6932 cells which passed quality filters per timepoint (Table S1). We clustered the cells using the Louvain algorithm, manually removed non-microglial clusters (which made up less than 3% of collected cells), and annotated the clusters *post hoc* (see Methods for details). We observed minimal expression of markers of *ex vivo* microglial activation^47^ (Fig. S2B).

**Fig. 2.**
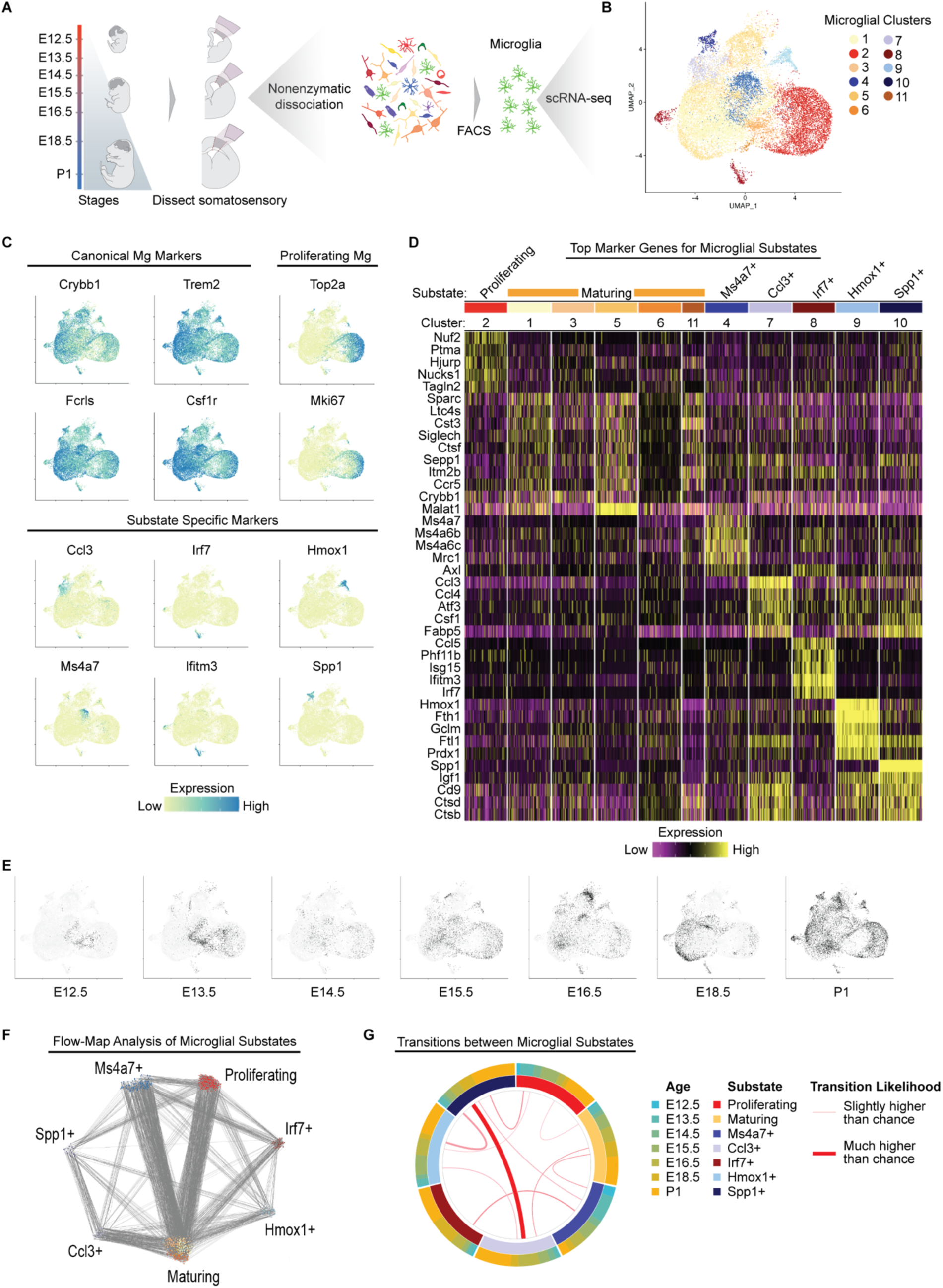
Comprehensive atlas of embryonic microglia. (A) Overview of experimental design for isolation of microglia for scRNA-seq. (B) UMAP visualization of microglia scRNAseq data from the developing somatosensory cortex of WT mice (combined UMAP representing 11 samples of 5-10 pooled embryos each, n= 21,135 total microglia). Cells are colored by microglia cluster assignment. (C) UMAP plots from B colored for expression of canonical, proliferating and substate specific microglial marker genes. (D) Heat map of top differentially-expressed genes in each microglia substate, grouped by cluster. (E) UMAP plots from B colored by time. (F) Force-directed weighted graph created by FLOWMAP algorithm representing microglia substate transitions. Cells are grouped by substates and each gray line represents one likely transition between cell types due to transcriptional similarity. The largest number of transitions occur between Ms4a7+ and Maturing microglia and between Proliferating and Maturing microglia. (G) Overrepresented microglial substate transitions across time created with BioCircos R package. The inner circle represents each substate, while the outer circle shows the distribution of ages of cells within each substate. Lines connecting substates depict transitions that happen more likely than chance when taking into account the number of cells within each substate as calculated by a Fisher’s exact test (FDR adjusted p<0.01). Line thickness represents the odds ratio of a transition as calculated by the Fisher’s exact test.

Our analysis revealed 11 microglial clusters present during embryonic corticogenesis (Fig. 2B). All clusters expressed canonical microglial markers such as *Crybb1* and *Trem2* (Fig. 2C). We did not find evidence of *Hoxb8* expression in any cluster at any timepoint^48^. Clusters 1, 3, 5, 6, and 11 were transcriptionally similar to one another, and expressed markers of embryonic microglia previously described in studies of normal microglial development and maturation^43,45^. Thus, these clusters were grouped together as likely representing a single substate, “maturing microglia.” The remaining clusters (2, 4, 7, 8, 9, and 10) expressed canonical microglial markers and also displayed high expression of identifying marker genes that were minimally expressed in other clusters; these clusters were annotated as unique microglial substates (Fig. 2C, D). Substate diversity increased over time, and substates with more unique gene expression profiles (clusters 7, 8, 9 and 10) displayed an overall higher relative abundance in later embryonic ages (Fig. 2E).

### A cluster of BAM-like microglia appears early in corticogenesis

Microglia cluster 4 expressed border-associated macrophage (BAM) markers (*Ccr1*, *Dab2*, *Mrc1*) and Ms4a family chemosensory genes, including *Ms4a6c*, *Ms6a6b* and *Ms4a7* (Fig. S3A, B). We therefore annotated this cluster as “*Ms4a7*-expressing microglia.” We confirmed that *Ms4a7*-expressing microglia are present in the embryonic cortical plate by multiplex RNA *in situ* hybridization (Fig. S3C). Microglia are derived from the same yolk-sac progenitors as BAMs, which appear in the meninges and vasculature around E10.5^49^. It is thought that local environmental signals trigger initiation of a transcriptional cascade that results in microglial identity after progenitor cells enter the brain parenchyma. Microglia that expresses BAM markers have been reported in the E14.5 mouse brain, but it is unknown when these cells first appear, how long they persist, or what the lineage relationship is between these cells and other microglia present in the embryonic and early postnatal brain^43^. In our dataset, cluster 4 microglia were present throughout embryonic development, even at the earliest age profiled (E12.5), prior to the emergence of most other microglial substates (Fig. 2E). To gain further understanding of the maturation and function of *Ms4a7*- expressing microglia throughout corticogenesis, we performed differential gene expression using DESeq2 with age treated as a continuous variable of interest, identifying genes with significant changes in expression over time within cluster 4 (Fig. S3D and Methods). GO analysis of the resulting up- and down-regulated genes revealed that cluster 4 microglia have more proliferative and metabolically active phenotype early in development, and later during corticogenesis, increase expression of genes involved in immune and neurodevelopmental functions (Fig. S3E, F).

### Embryonic microglial proliferation has two peaks

Microglia cluster 2 was characterized by high expression of markers of active cell cycling, such as *Ube2c*, *Hist1h1b*, and *Mki67*, and was therefore annotated as “Proliferating microglia” (Fig. S4A, B). Cluster 2 peaked in overall proportion of microglia early during neurogenesis, at approximately E14.5, decreased at intermediate ages, and increased again at P1, suggesting that birth may act as an immune stimulus for microglial proliferation (Fig. 2E). Using RNA *in situ* hybridization, we confirmed the presence of proliferating, *Ube2c*-expressing microglia located in the dorsal cortical ventricular zone, corresponding to the previously described localization of proliferating microglia^50^ (Fig. S4C). Proliferating (cluster 2) microglia expressed BAM markers such as *Ms4a6c* and *Pf4* at early developmental ages only (E12.5-E13.5). At later ages, cluster 2 microglia showed increased expression of markers of cytokine production and immune cell differentiation and activation (Fig. S4D-F). Thus, over the course of embryonic development, proliferating microglia may also take on more of the immune functions classically attributed to microglia. Alternatively, maturing microglia or other subspecialized substates may revert to a proliferative state during later stages of corticogenesis, while retaining aspects of a more differentiated transcriptional signature.

### Microglial substate diversity increases throughout embryonic development

We observed four clusters (clusters 7, 8, 9 and 10) that did not express BAM markers and that increased in proportion in the late embryonic period (Fig. 2E). These clusters displayed high expression of genes and gene modules unique to each substate (Fig. 2D). Cluster 10 was characterized by high expression of osteopontin (*Spp1*), a pro-inflammatory phosphoglycoprotein that is secreted by microglia under stress conditions, such as in stroke, neurodegenerative disorders, and multiple sclerosis ^51^ (Fig. 2D). The gene expression profile of cluster 10 resembled the expression profile of axon tract-associated microglia previously described in the corpus callosum and cerebellar white matter of the P4/5 mouse brain^43^. Cluster 7 shared several marker genes with cluster 10 but lacked high *Spp1* expression and displayed upregulation of distinct pro-inflammatory cytokines, including *Ccl3* and *Ccl4* (Fig. 2D). Cluster 9 was characterized by high expression of heme oxygenase (*Hmox1*), an enzyme that catalyzes the rate-limiting step of heme degradation (Fig. 2D). Microglial *Hmox1* is thought to reduce inflammation after lipopolysaccharide (LPS) exposure and to improve neurorecovery after intraventricular hemorrhage, but the role of microglial *Hmox1* in normal brain development is unknown^52,53^. Cluster 8 microglia displayed upregulation of several interferon-response genes, including *Irf7*, *Ifitm3*, and *Ifit3* (Fig. 2D). These genes are involved in regulation of the inflammatory response and inhibition of the viral life cycle. The transcriptional profiles of cluster 8 and 7 microglia resembled those of specific populations of *Ifitm3*+ and *Ccl4*+ microglia, respectively, that have previously been reported in the aged brain^43^.

Given that clusters 7, 8, 9 and 10 appeared later in developmental time, lacked BAM marker expression and expressed markers of mature microglia, we hypothesized that *Ms4a7*-expressing microglia may precede these later substates in a lineage relationship. We analyzed our scRNA-seq data using FLOW-MAP, a graph-based algorithm for trajectory mapping that incorporates sequential timepoint information ^54^ (Fig. 2F). FLOW-MAP analysis suggested that *Ms4a7*-expressing microglia are most likely to have a direct lineage relationship with maturing microglia and with *Irf7*- and *Spp1*-expressing substates (Fig. 2G). *Hmox*- and *Ccl3*-expressing microglia are most likely to transition directly from *Spp1*-expressing microglia (Fig. 2G). FLOW-MAP analysis did not suggest a consistent, unidirectional lineage pathway between microglial substates, likely reflecting the dynamic nature of microglial activation, proliferation, and dormancy in the presence of variable external stimuli. Overall, these data constitute a comprehensive map of microglial substates throughout corticogenesis, with high temporal and cell type resolution, and will serve as a resource for future studies of microglial function in both normal development and in disease states.

### All microglial substates highly express receptors for TORCH pathogens and cytokines across development

To understand whether specific microglial substates are poised to respond to inflammatory and infectious insults during brain development, we quantified expression over time of TORCH pathogen receptors and cytokine receptors using our single-cell transcriptional atlas of developing microglia (Fig. 3A). We observed that all microglial clusters displayed relatively high, sustained expression of multiple receptor types from E12.5 through P1, and all clusters had high CytoTORCH module scores (Fig. 3A, B). We compared expression of specific CytoTORCH receptors in microglia and other cortical cell types using the single-cell atlas of cortical development employed above^22^. Some receptors, such as *Tlr2* and *Tlr12*, demonstrated negligible expression in neurons and neural progenitors, but were highly expressed in multiple microglial substates, suggesting a possible primary role for microglia in mediating the corresponding infections (Fig. 1A, 3A). TLR2 is a receptor for *Treponema pallidum*, which causes congenital syphilis. This disease is associated with many neurological problems, including microcephaly, ventriculomegaly, seizures, developmental delays, and intellectual disability^55^. *Tlr2* showed a gradual increase in expression over embryonic development in most microglial substates, and no significant expression in other embryonic cortical cell types. TLR12 functions as a receptor for *Toxoplasma gondii*, the parasite that causes congenital toxoplasmosis, which is associated with intracranial calcifications, hydrocephalus, seizures, developmental delay, and learning disabilities^55^. *Tlr12* expression was highest in proliferating microglia (cluster 2), maturing microglia (cluster 1), and Ms4a7+ microglia (cluster 4). *Tlr12* was also highly expressed in pericytes and endothelial cells (Fig. 1A), suggesting that these cells may help mediate CNS entry. Other receptors displayed high expression across many cell types in the embryonic mouse brain. For instance, *Axl*, a receptor for Zika virus, was highly expressed in most microglial substates (Fig. 3A), in pericytes and endothelial cells, and in many cells early in the excitatory neuronal lineage, including APs and IPs (Fig. 1A). These patterns of receptor expression suggest that some pathogens can directly target multiple cell types in the fetal brain parenchyma, while others initially act on microglia, which may in turn affect surrounding cell types.

**Fig. 3.**
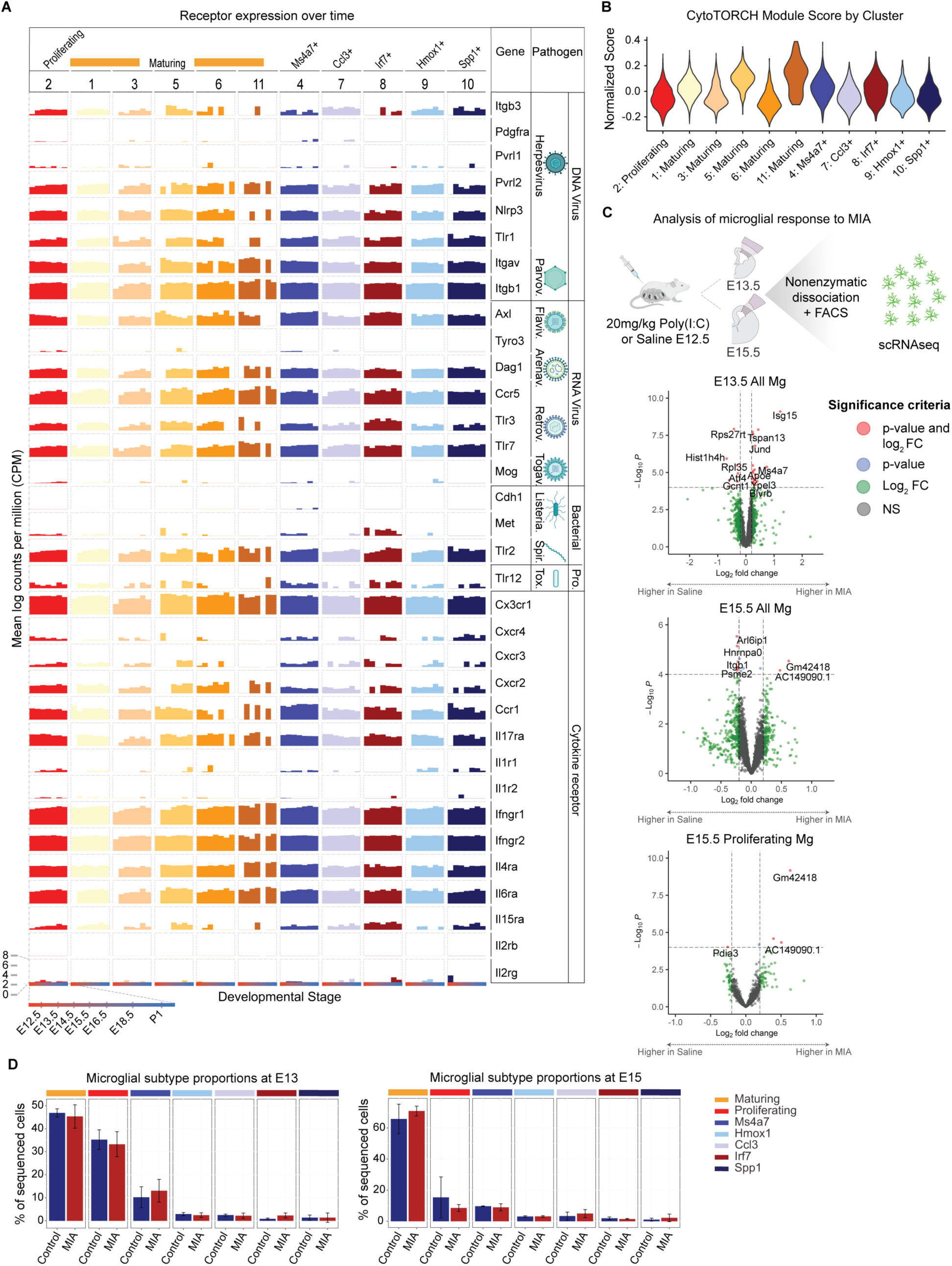
Microglial substates transcriptional changes after MIA. (A) Bar plot of TORCH pathogen and cytokine receptor expression (mean log counts per million) in each microglia cluster across development using developmental microglial atlas. (B) Violin plot of CytoTORCH module score for each microglia cluster. (C) Overview of experimental design (up) and volcano plots (down) showing differentially expressed genes in microglia from the embryonic prospective somatosensory cortex after maternal Poly(I:C) or saline injection. A total of 30,985 cells were sequenced, comprising 6 pooled samples from a total of 37 embryos for saline-treated controls, and 6 pooled samples from a total of 32 embryos for MIA. Genes are color coded according to enrichment (log2FC > 0.15 and −log10Benjamini-Hochberg adj. P<0.05). Dotted lines represent enrichment cut-offs. Top signature genes are annotated. Log foldchange and p-values were generated with a generalized linear model with Benjamini-Hochberg correction using DESeq2. (D) Bar plot showing embryonic microglial substate proportions after maternal injection of saline as compared to Poly(I:C). Generalized mixed effects binomial models showed no significant changes in substate proportions between treatment groups. Error bars represent 1 standard deviation in substate proportion. Illustrations created with BioRender.com.

Microglia displayed higher and more sustained expression over time of receptors for systemic inflammatory stimuli and Poly(I:C) than most other cell types in the developing brain (p<0.05 for ANOVA test and for pairwise comparisons between microglia and all other cell types using Tukey’s HST). *Tlr3*, which encodes the receptor for Poly(I:C), was expressed throughout development in all microglia except for one maturing microglia cluster (cluster 11). *Tlr3* was moderately expressed in astrocytes, endothelial cells, and pericytes, primarily in late embryonic development, and minimally expressed in neurons and neural progenitors. All microglial substates displayed high, sustained expression of receptors for cytokines known to be involved in the response to MIA, including *Il17ra* and *Il6ra*. All microglial substates also highly expressed receptors for a subset of cytokines involved in maternal fever, including IL-6, IL-8, and interferon-gamma^56,57^. These receptor patterns suggest that microglia, as compared to most other cell types in the dorsal cortex, are more poised to respond to MIA and systemic inflammatory stimuli during fetal development.

### Single-cell transcriptional changes in microglia in response to maternal immune activation

We asked how embryonic microglia respond to an *in utero* inflammatory stimulus at the single-cell level, hypothesizing that MIA leads to upregulation of genes involved in the inflammatory response. We performed scRNA-seq of FACS-isolated microglia from the prospective somatosensory cortex at E13.5 and E15.5, after MIA or sham saline injection at E12.5 (Fig. 3C). In total, we collected 35,794 scRNA-seq profiles, with 12,046 to 23,748 cells per timepoint. We clustered the cells as described above, manually removed non-microglial clusters, and annotated the clusters *post hoc* (Methods and Fig. S5A). This resulted in 30,985 cells in 11 microglial clusters suggestive of 7 unique substates that closely mirrored the clusters from our analysis of normal embryonic microglial development (Fig. S5B). We found no changes in microglial substate proportions at E13.5 or E15.5 after MIA using a binomial generalized linear mixed effects model (Fig. 3D).

DEGs at each timepoint in all microglia and in each substate between MIA and saline control were identified by DESeq2 using pseudobulked counts and a negative binomial generalized linear model. We first assessed MIA-induced single-cell gene expression changes at E13.5, 24h after the immune stimulus (Fig. 3C). When analyzing all E13.5 microglia together, we observed an upregulation of pro-inflammatory genes including *Ccl12*, *Jund*, *Isg15*, and *Clec12a* after MIA. GO analysis (see Methods for details) also demonstrated a significant enrichment in genes involved in “Response to Interferon-Beta” (Fig. S6A). MIA led to a reduction in expression of genes involved in proliferation when all microglia were grouped and analyzed together, with associated GO terms including “Chromatin organization” and “Gene silencing” (Fig. S6B). Analyzing the response to MIA in each cluster separately, we detected significant DEGs in maturing, proliferating, and Ms4a7+ microglial substates, but not in other substates (Table S3). This result was likely related to the small number of microglia in many embryonic substates (Fig. 2E). Maturing microglia displayed the largest number of upregulated genes, many of which were associated with GO terms related to energy metabolism and immune activation, such as “Positive regulation of leukocyte migration” and “Myeloid dendritic cell activation” (Fig. S6A). Overall, 24h after an immune stimulus, most microglia in the embryonic dorsal cortex displayed gene expression changes suggestive of increased activation and chemotaxis, and we did not find evidence of a specific, selectively activated microglial substate.

We next analyzed single-cell gene expression changes in microglia at E15.5, 3d after MIA (Fig. 3C). The widespread upregulation of genes involved in microglial activation that had been present at E13.5 did not remain at E15.5 (Table S4). Significantly enriched GO terms for upregulated genes included “Aging” and “ATP synthesis coupled electron transport” (in proliferating microglia). Significantly enriched GO terms for downregulated genes included “Response to endoplasmic reticulum stress” (maturing microglia) and “Proteasomal ubiquitin-independent protein catabolic process” (maturing microglia) (Fig. S6B). Overall, gene expression changes in microglia 72 hours after MIA were suggestive of persistent alterations in energetics and protein catabolism, mirroring effects seen at E15.5 in non-microglial cell types (Fig. 1A, B).

### MIA-exposed microglia display persistent gene expression changes in juvenile mice

Prenatal MIA alters social behaviors in adolescents and was recently demonstrated to impair the microglial response to immune challenges in adult animals^13,21^. We hypothesized that persistent alterations in gene expression would be detectable in microglia from juvenile mice exposed to MIA during embryonic development. We performed scRNA-seq of FACS-isolated microglia from P14 somatosensory cortex after MIA or sham saline injection at E12.5 (Fig. 4A). After quality control and doublet filtering (see Methods and Table S1), we analyzed the expression profiles of 28,474 cells. We clustered the cells using the Louvain algorithm, manually removed non-microglial clusters^46,58^, and annotated the clusters *post hoc* (Fig. 4A). We observed seven clusters of microglia that comprised five distinct substates (Fig. 4B). Cluster 1, 2 and 3 microglia expressed canonical microglial markers and homeostatic genes and were thus annotated as a single substate, “homeostatic microglia.” The other clusters expressed subsets of non-homeostatic genes and were annotated as additional microglial substates. Cluster 4, “Apoe+ microglia,” was characterized by high *Apoe* expression. Cluster 5, “inflammatory microglia,” expressed pro-inflammatory cytokines including *Ccl3* and *Ccl4*, and the cytokine receptors *Cd9* and *Cd63*, which are also associated with AD plaques^59^. Cluster 6, “cycling microglia,” was composed of microglia expressing genes involved in proliferation and the cell cycle, while cluster 7, “innate immune microglia,” had high expression of genes involved in the innate immune response, including interferon-responsive genes such as *Ifitm3* and *Bst2*. Microglial substate proportions did not differ significantly at P14 between MIA and saline injection (Fig. 4C).

**Fig. 4.**
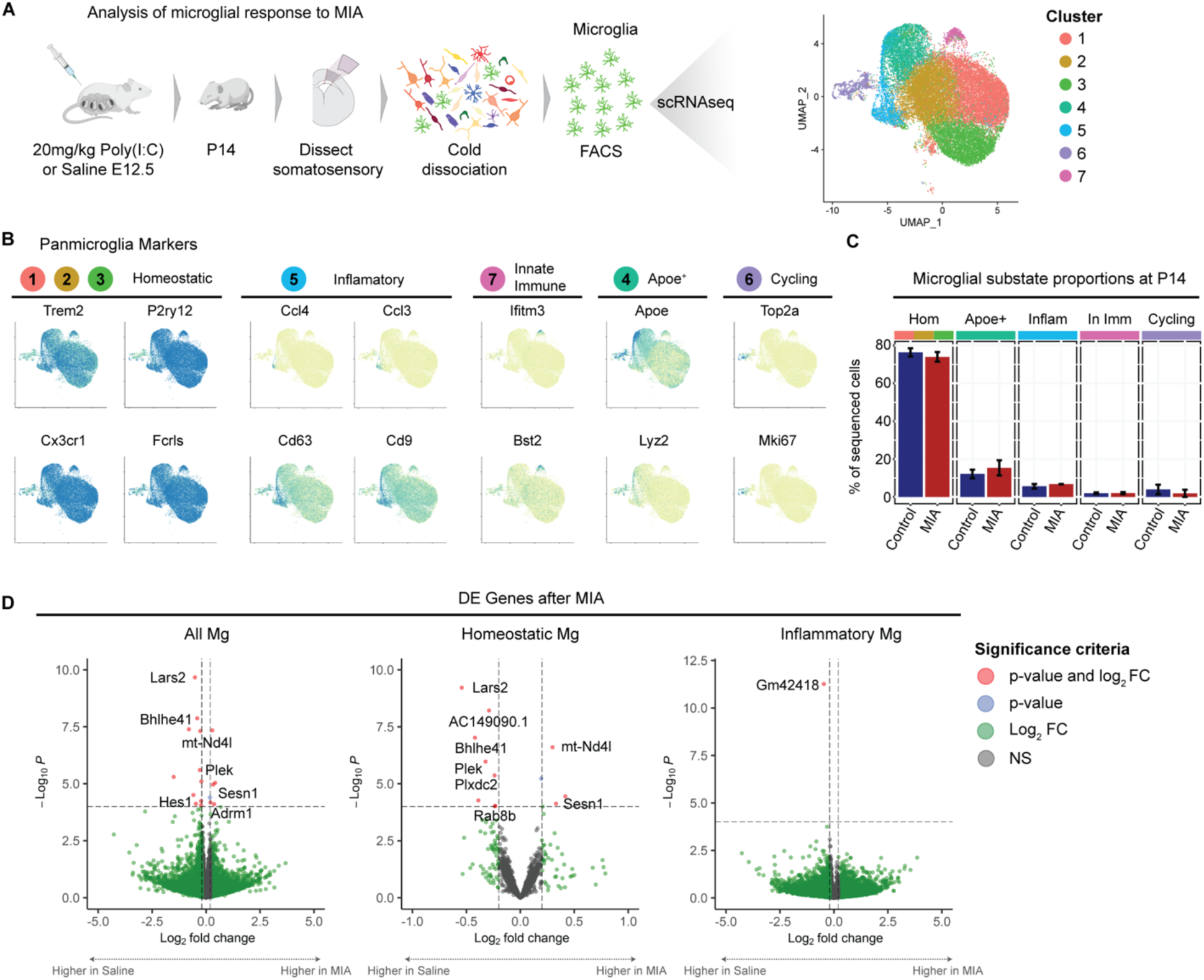
Persistent transcriptional changes in microglia after MIA. (A) Overview of experimental design and UMAP visualization of microglia scRNA-seq data from the somatosensory cortex of P14 mice after maternal injection of saline or Poly(I:C) at E12.5. In total, we sequenced 28,474 microglia from a total of 6 P14 mice, 2 of which were treated with Poly(I:C) and 4 of which were treated with saline. Cells are colored by cluster. (B) UMAP plots from A colored for expression of homeostatic, inflammatory, innate immune, Apoe+ and cycling substate specific microglial marker genes. (C) Bar plot showing P14 microglial substate proportions after maternal injection of saline as compared to Poly(I:C). Generalized mixed effects binomial models showed no significant changes in substate proportions between treatment groups. Error bars represent 1 standard deviation in substate proportion. (D) Volcano plots showing differentially expressed genes in microglia from the P14 somatosensory cortex after maternal Poly(I:C) or saline injection. Genes are color coded according to enrichment (log2FC > 0.15 and −log10Benjamini-Hochberg adj. P<0.05). Dotted lines represent enrichment cut-offs. Top signature genes are annotated. Log foldchange and p-values were generated with a Generalized linear model with Benjamini-Hochberg correction using DESeq2. Illustrations created with BioRender.com.

DEGs between MIA and saline control injection were assessed in each cluster and in all microglia grouped together using DESeq2 and pseudobulked counts (see Methods for more details). At P14, a small subset of genes were significantly differentially expressed in MIA-exposed as compared to saline-exposed microglia. Significantly downregulated genes include *Plek* and *Bhlhe4*, which are known to regulate microglial phagocytosis, and *Hes1* and *Plexdc2*, which promote M2 polarization^60–63^ (Fig. 4D and Table S5). Thus, after exposure to an inflammatory stimulus *in utero*, microglia in juvenile animals may have an altered capacity for activation and phagocytosis.

## DISCUSSION

Much remains unknown regarding how maternal inflammation during pregnancy leads to adverse neuropsychiatric outcomes. Our findings demonstrate that microglia are necessary for the response of many embryonic cortical cell types to *in utero* inflammation, including neurons and neuronal progenitors. Microglia can play opposing roles in different types of brain pathology, sometimes helping promote recovery and pathogen clearance, while other times exacerbating inflammation and enabling viral replication^16,19,64,65^. When considering microglia as a potential future therapeutic target, it will be essential to determine whether the downstream effects of microglial signaling in the setting of maternal inflammation are beneficial or harmful to the developing brain.

Our atlas of developing microglial gene transcription suggested that most microglia in the embryonic brain are capable of responding to a variety of infectious and inflammatory stimuli over a prolonged time window. High expression of CytoTORCH receptors was also observed in pericytes and endothelial cells, reflecting the important role of these cell types in pathogen entry into the CNS. Despite relatively uniform expression of CytoTORCH receptors across microglia, many substates bore distinct transcriptional signatures, and we observed rapid changes in substate proportions at high temporal resolution. *Ms4a7*-expressing microglia were present at the youngest ages studied, and FLOW-MAP analysis was consistent with this substate possibly representing an early link in the lineage relationship between yolk sac progenitors and other microglial substates. Four substates (clusters 7, 8, 9 and 10) appeared around E15.5 and increased in proportion thereafter, suggesting the potential for increased subspecialization of microglial function in later stages of corticogenesis. Three of these later-appearing clusters (7, 8, and 10) displayed gene expression profiles similar to those of microglia in the aged or diseased brain^43,66^. These results suggest that developmental gene transcriptional programs could become reactivated in microglia later in life in response to environmental or genetic factors. Future studies using genetic labeling and conditional knockout methods are needed to unravel substate-specific roles in development and disease, and to confirm the lineage relationships between *Ms4a7*-expressing microglia and other substates.

We queried the temporal dynamics of the microglial response to *in utero* inflammation, observing that embryonic microglia upregulated pro-inflammatory and interferon-response genes 24 hours after an immune stimulus. These effects mirror microglial profiles previously described *in vitro* after Poly(I:C) or LPS stimulation, and *in vivo* in the setting of “pseudo-TORCH” syndromes, multiple sclerosis, and brain ischemia^67,68^. The immediate pro-inflammatory response subsided by E15.5, and was followed by more subtle, sustained changes in microglial gene expression that persisted through P14. Additional studies are needed to understand the implications of these transcriptional changes. Recently, it has been demonstrated that microglia exposed to MIA during development display an altered ability to respond to inflammatory stimuli in adulthood^21^, suggesting that the transcriptional changes we observed at P14 may reflect a persistent protective state that could reduce the harmful effects on the young brain of chronic microglial activation due to recurrent inflammatory stimuli.

Taken together, these findings confirm the remarkable heterogeneity of microglia during development and show that microglia are essential for the response of other cell types in the fetal brain to *in utero* inflammation. A mechanistic understanding of how such responses are established may inform future therapeutic strategies to reduce the chances of adverse neuropsychiatric outcomes in fetuses exposed to maternal infections or inflammatory insults.

## DATA AVAILABILITY STATEMENT

All data are available upon reasonable request from Lead Contact.

## ACKNOWLEDGMENTS

We thank members of the Arlotta and Levin laboratories for scientific input, technical assistance and editing of the manuscript. We thank Juliana Brown for assistance with writing the manuscript and editing the figures and Sara Simes and Vy Vuong for assistance with mouse genotyping and brain tissue sectioning. We thank the Bauer Core Facility and Flow Cytometry Core at Harvard and the Harvard Center for Biological Imaging for their important contributions to this work.

## AUTHOR CONTRIBUTIONS

B.E.L.O., J. S. and P.A. conceived the project. B.E.L.O., N.D.I., and J.S. performed the majority of the experiments with guidance and support from P.A. Bioinformatics and scRNAseq analysis was performed by B.E.L.O., J.S., T.F., and K.K., with guidance and support from P.A. and J.Z.L. B.E.L.O., N.D.I. and P.A. co-wrote the paper with input from all authors. All authors approved the final version for submission.

## METHODS

### EXPERIMENTAL MODEL AND SUBJECT DETAILS

#### Mouse Lines

WT mice – Developing microglia atlas and microglial transcriptional response to MIA experiments were performed using C57Bl/6N WT mice from Charles River (strain code 027).

*Csf1r* Knockout – The *Csf1r* KO mouse line was procured from Jackson Laboratories (stock #028064) and was maintained on a C57Bl/6N background from Charles River (strain code 027).

#### Genotyping

*Csf1r* mice were genotyped using the following primers: CSF1R forward primer: TCCCAGCATTAGGCAGCCT; CSF1R reverse primer: GCCACCATGTGTCCGTGCTT. As we used male mice only for snRNAseq experiments using the *Csf1r* KO mouse line, sex was also determined for embryonic mice via amplification of the *Sry* gene on chromosome Y using the following primers: Sry forward primer: ACAAGTTGGCCCAGCAGAAT; Sry reverse primer: GGGATATCAACAGGCTGCCA. Sex of WT embryos for scRNAseq experiments (microglial atlas and microglial response to MIA) was not distinguished.

#### Mouse husbandry

All animal procedures were approved by the Harvard University IACUC and conducted in compliance with institutional and federal guidelines. Mice were housed in individually ventilated cages using a 12-hour light/dark cycle and had access to water and food *ad libitum*. Males only were used for snRNA-seq experiments of cortex from *CSF1R* mutant versus WT mice, given the published male predominance of behavioral effects and changes in neuronal gene expression after MIA^12,32,33^. Equal ratios of male and female mice were used in scRNA-seq of FACS-purified microglia in postnatal samples in which sex could be determined. We also used both female and male mice in scRNA-seq experiments of FACS-purified embryonic microglia but were unable to prospectively determine sex to verify equal ratios of males and females. This limitation existed because tissue was processed for single cell isolation and sequencing immediately following brain dissection to minimize *ex vivo* microglia activation, which did not allow time for genotyping between embryo dissection and sample preparation.

## METHOD DETAILS

### Maternal Immune Activation

Maternal immune activation was induced by intraperitoneal (IP) injection of pregnant dams at E12.5 with poly(I:C) as previously described^69^. Briefly, C57Bl/6 WT timed pregnant female mice or CSF1R^+/-^ female mice that had undergone timed mating with CSF1R^+/-^ male mice were injected intraperitoneally at E12.5 with 20mg/kg Poly(I:C) (Sigma, P0913) resuspended in 0.9% sterile sodium chloride (Thermo Fisher Scientific NC9016275). Control animals were injected with an equal volume of 0.9% sodium chloride. Confirmation of the expected maternal inflammatory response using this protocol was done by cytokine analysis via a membrane-based immunoassay per manufacturer instructions (R&D Systems, ARY006), performed on maternal splenic tissue obtained after sacrificing animals not used for sequencing experiments 5 hours after Poly(I:C) injection. Cytokine levels were assessed using the Image Lab software (BioRad) and statistics performed with GraphPad Prism (t-test with Holm-Sidak multiple comparison correction). Mice were note disturbed after MIA until the time of euthanasia for RNA analysis.

### RNA Fluorescent *in situ* Hybridization and Immunohistochemistry

Brains for ISH were prepared by transcardial perfusion of ice-cold Phosphate Buffered Saline (PBS), then ice-cold 4% paraformaldehyde (Electron Microscopy Sciences, 15710) diluted in PBS. Brains were stored overnight in 4% paraformaldehyde at 4°C, followed by sequential washes with PBS and gradually increasing concentrations of sucrose in PBS at 4°C for cryoprotection per the ACDBio Multiplex Fluorescent V2 Assay kit protocol (ACDBio, 323100). Brains were frozen in a solution of 2-parts 30% sucrose, 1-part OCT (Tissue-Tek, 4583) in Peel-A-Way embedding molds (VWR, 15160-215) and stored at −80°C. PFA-fixed tissues were cryosectioned at 16 um. For ISH tissue was heated at 70°C prior to use to adhere tissues to glass slides. Sections were hydrated with PBS for 5 minutes, dried and re-hydrated 5 minutes to remove residual OCT. Target retrieval solution (ACDBio 322000) was heated to a rolling boil and incubated on the slides for 5 minutes followed by two quick rinses of water and dehydration with 100% ethanol. Protease III (<P14 tissues) or Protease IV (P60 tissues) reagent (ACDBio, 322340) was applied to the slides for 15-60 minutes, depending on age of the tissue, at 40°C. Slides were washed 2x in water, followed by pooled probe hybridization and signal amplification according to manufacturer’s protocols. RNAscope Catalog Target Probes used were: *C1qa* (ACDBio, 441221-C3), *Ube2c* (ACDBio, 552191-C2), and *Ms4a7* (ACDBio, 314601). For immunohistochemistry, tissue was blocked and permeabilized with for 2 hours at room temperature in blocking solution (PBS with 0.5% BSA, 0.3% Triton X-100, and 2% donkey serum), and incubated overnight at 4°C in blocking solution with rabbit anti-Iba1 (1:100, Wako, 019-19741). After washing of primary antibody, secondary antibody incubation was done for 2h at room temperature in blocking solution with donkey anti-rabbit IgG (H+L) Alexa Fluor 555.

### Imaging

Confocal imaging of RNAscope *in situ* hybridization-stained tissue sections was performed using a Zeiss 700 inverted confocal microscope. Tile scan images spanning the width of the dorsal cortical wall, from pia to corpus callosum, were captured using a 20x air objective and stitched using Zeiss Zen Software (version 2.6). Z-stack images were acquired using the 63x oil-immersion objective with a 0.5 um Z-step. All confocal imaging was captured using Zen Black software (Zeiss, v2.3). Fluorescence imaging of Iba1 immunostaining was performed using a Zeiss Axio Imager 2 upright microscope with a 20x air objective using Zeiss Zen Software.

### Tissue Dissection for RNA Sequencing

For experiments involving snRNA-seq or scRNA-seq of embryonic brain tissue, we euthanized the pregnant females at the age of analysis and obtained the embryos. For scRNA-seq of postnatal microglia, mice were housed with their mothers and littermates until euthanasia on the day of analysis. Brain dissection was performed with RNAse-free technique in ice cold Hybernate-E (Brainbits) for embryonic tissue or in ice cold Hibernate-A (Brainbits) for postnatal tissue. The somatosensory (S1) cortex (or prospective S1 cortex for embryonic tissue) was dissected and meninges removed. Tissue was immediately processed as described below for scRNA-seq experiments or flash frozen in liquid nitrogen and stored at −80°C until genotypic was completed for snRNA-seq experiments.

### Single-cell Suspension Preparation and FACS

Live microglial suspensions from dissected tissues were prepared for fluorescence activated cell sorting (FACS) and maintained under ice-cold conditions as previously described^46^. Prospective S1 cortex blocks were minced and transferred to a 5 mL microcentrifuge tube with Hibernate-A and pelleted at 500 *g* for 5 min, and then the supernatant was decanted. The pellet was resuspended in 1 mL dissociation buffer (20 mL PBS (Gibco, 15630-080) + 20 uL RNasin (Promega, 40 units/ul, N2515) + 400 uL DNAse (Worthington, 12,500 units/mL, LS002006)) and transferred to a 2 mL Dounce homogenizer (Sigma D8938-1SET). The tissue was homogenized with 15 plunges of pestle “A”. Homogenate was passed through a 70 um cell strainer (Miltenyi, 130-110-916), then pelleted at 500 *g* for 5 min. The pellet was resuspended in 500 uL FACS buffer (PBS with 0.04% Bovine Serum Albumin (BSA, NEB, B9000S)). A combination of three primary antibodies, anti-Cd11b-PE (BioLegend, 101208), anti-Cd45-APC/Cy7 (BioLegend, 103116), and anti-Cx3cr1-APC (BioLegend 149008) were diluted 1:200 in 500 uL FACS buffer and incubated with the cells at 4°C for 15 minutes. The labelled cells were washed with 5 mL PBS, pelleted at 500 *g* for 5 min, resuspended in 500 uL of FACS buffer, and finally passed through a 40 um filter (Corning, 352235).

For the microglial atlas and embryonic microglial MIA experiments, prospective S1 cortical dissections from 2-8 embryos per age per experiment were pooled, homogenized, sorted, and processed for scRNA-seq. At least two biological replicates for each age range (E12.5/13.5 [birthdate of layer 5 and 6 excitatory neurons], E14.5-E16.5 [birthdate of layer 2/3 and 4 excitatory neurons], E18.5-P1 [end of neurogenesis, transition to gliogenesis]) were obtained for the microglial atlas experiments. At least three biological replicates (pooled sets of embryos) per treatment group within each age were obtained for microglial MIA experiments. For the postnatal MIA experiments, S1 cortical blocks were dissected and pooled, homogenized, sorted, and processed for scRNA-seq. Two biological replicates (pooled sets of 1 male and 1 female) were obtained for the Poly(I:C)-treated condition and four biological replicates (two pooled sets of 1 male and 1 female, one samples from an individual male and sample from an individual female) were obtained for the saline-treated condition.

### Single-nucleus Suspension Preparation and FACS

Prospective S1 cortices of E15.5 embryos after MIA at E12.5 were dissected and flash-frozen in liquid nitrogen and stored at −80°C. Tail samples were simultaneously obtained from each embryo followed by DNA extraction and genotyping (see genotyping methods above) for sex and *Csf1r* genotype. Frozen tissue blocks from male *Csf1r* mutant and WT embryos were processed to obtain a single-cell suspension using papain digestion (15 minutes) (Papain dissociation kit, Worthington), following the manufacturer’s protocol. After dissociation and concentration, the pellet was resuspended in 500 uL of Nuclei Solubilization Buffer (10 mL PBS (Gibco, 15630-080) + 50 uL of 20 mg/mL BSA (NEB, B9000S) + 10 uL RNasin (Promega, N26515)) spiked with 1 uL of Hoechst and passed through a 40 um filter (Corning 352235). Nuclei were sorted on a Beckman Coulter MoFlo Astrios EQ Cell Sorter, pre-chilled to 4°C, using a 100 um nozzle, and collected into wells of a 96-well plate pre-filled with 10 uL of Nuclei Solubilization Buffer, and immediately processed for single-cell GEM formation (10x Genomics, single cell RNA sequencing 3’, Chromium v3.1). We obtained 3 biological replicates of 1-2 pooled embryos per treatment condition.

### Single-cell RNA Sequencing (scRNA-seq) and Single-nucleus RNA Sequencing (snRNA-seq)

scRNA-seq libraries were prepared with the Chromium Single Cell 3’ Kit v3.1 (10x Genomics, PN-1000121). Sorted live, dissected microglia suspended in FACS buffer or sorted nuclei (15,000 nuclei/sample) suspended in Nuclei Solubilization Buffer were loaded onto a v3.1 Chromium Single Cell 3’ G Chip (10x Genomics, PN-1000127) and processed through the Chromium controller. scRNA-seq libraries from different dissected samples were pooled based on molar concentrations and expected captured cell counts, then sequenced on a NovaSeq (Illumina) instrument with 28 bases for read 1, and 91 bases for read 2. The mean reads per cell in each sample was 35K. snRNA-seq libraries were sequenced on a NovaSeq (Illumina) instrument with 26 bases for read 1, 91 bases for read 2, and 8 bases for the i7 index. The mean reads per nucleus was >24K with a minimum sequencing saturation of >40.5%.

### Single nucleus RNA-seq data pre-processing and clustering

We performed snRNA-seq using the 10x Chromium platform (v3.1) on eight male mouse samples separately, extracted at E15.5, split into four experimental groups: two *Csf1r* null samples with MIA (each sample from one embryo), two *Csf1r* null samples with saline injection (one sample from one embryo, one sample from two pooled embryos), two WT samples with MIA (each sample from one embryo), and two WT samples with saline injection (one sample from one embryo, one sample from two pooled embryos). We aligned raw sequences to mm10 and produced cell-by-gene count matrices for each sample using 10X Genomics Cell Ranger 6.0.1. We used Seurat v4.0.0 to remove cells with fewer than 200 expressed genes or fewer than 500 UMIs. We scaled each sample independently before integrating all samples into a single Seurat object using Seurat’s SelectIntegrationFeatures(), FindIntegrationAnchors(), and IntegrateData() functions.

We performed principal component analysis (PCA) on the 2000 genes used for dataset integration, and used the top 30 components for Shared Nearest Neighbor (SNN) Louvain clustering with a resolution parameter of 0.4, as well as for Uniform Manifold Approximation and Projection (UMAP) for visualization. We found 17 clusters, and manually annotated the cell type identity of each cluster based on marker genes found using Seurat’s FindAllMarkers() function. For downstream analysis, we collapsed clusters which shared cell type labels. Clusters with fewer than 200 cells across all samples were ignored in differential gene expression analyses and GO term enrichment between experimental groups.

### Differential expression analysis and GO enrichment

We used NEBULA to perform pairwise differential expression analyses within each cell type between *Csf1r* null mice with MIA vs saline control, as well as between WT mice with MIA vs saline control. We used sample identity as a random effect, MIA treatment vs control as a fixed effect, and UMI count per cell as the offset. We performed Benjamini-Hochberg (BH) p-value adjustment and set a false discovery rate (FDR) threshold of 0.05. We achieved similar results when analyzing differentially expressed genes using DESeq2 and pseudobulked counts. For each cell type, genes were split into several lists based on whether they were significantly upregulated, significantly downregulated, or not significantly altered in response to MIA in *Csf1r* null mice and in WT mice according to our NEBULA analyses. We used the ClusterProfiler R package to perform GO term enrichment analyses on each of these gene lists. For lists with significantly enriched GO terms, we collapsed redundant and closely related GO terms using ClusterProfiler’s simplify() function with a similarity cutoff of 0.4.

### Purified microglia scRNA-seq data pre-processing and clustering

We aligned raw sequencing reads from all purified microglia scRNA-seq datasets against mm10 and produced cell-by-gene count matrices using Cell Ranger v6.0.0. We collected and processed 11 samples from untreated WT mice: 2 samples from each of timepoints E13.5, E14.5, E15.5, and P1, and a single sample from each of timepoints E12.5, E16.5, and E18.5. We merged these 11 samples into a single Seurat object and normalized them using SCTransform() before performing PCA, and then UMAP on the top 30 components. We then performed SNN Louvain clustering with a resolution parameter of 0.3 on the top 30 components, yielding 12 clusters. One of the clusters showed high expression of Sox11 and other neuronal markers and lacked expression of microglia markers, so we excluded it from downstream analysis as neuronal contamination, resulting in the final 11 clusters. Additionally, we collected and processed 12 embryonic microglial scRNA-seq samples after treating pregnant dams treated with Poly(I:C) (MIA) or saline injections (see Table S1 for number of pooled embryos collected for each age and condition) and 6 samples from pups at P14 (see Table S1 for number of pups per sex per condition). We removed cells with fewer than 500 expressed genes or a mitochondrial gene content higher than 20%. We merged all 12 embryonic samples into a single Seurat object and all postnatal samples into a second object using merge(), then normalized them using SCTransform(). We clustered cells using the SNN Louvain algorithm on the top 30 principal components, using a resolution of 0.5 for the embryonic cells and 0.8 for the postnatal cells. We removed one neuronal cluster from the embryonic dataset and two neuronal clusters from the postnatal dataset as contamination. We manually annotated the remaining clusters as microglial substates based on marker genes derived for each cluster using Seurat’s FindAllMarkers() function. We derived cell cycle scores using Seurat’s CellCycleScoring() function. Because our MIA vs saline purified microglial samples each included cells from both males and females, and because we observed sexual dimorphism in microglial expression profiles, we used *Xist* and *Tsix* expression to identify the most likely sex of individual cells.

### Expression changes over time in developing microglia

To assess changes in expression over the course of development within each microglial substate, we summed counts from all cells from each sample within each substrate to create a pseudobulked counts matrix. We assigned a numeric age value to each developmental timepoint corresponding to the number of days since E12.5 (eg, E12.5=0, E18.5=6, etc). We then used DESeq2 to perform differential expression analyses with age as a continuous variable.

### Differential expression in microglia in response to immune activation

We summed all counts for female (Xist+ and/or Tsix+) and male cells of each sample within each microglial substate to derive pseudobulk expression values. We passed those pseudobulked count matrices to DESeq2 to perform differential expression analyses between MIA and control independently for each substate at each timepoint, using sex and sequencing batch as covariates. For each microglial subtype and timepoint, we performed GO term enrichment using ClusterProfiler() on the lists of significantly (FDR < 0.1) upregulated and downregulated genes. For lists with significantly enriched GO terms, we collapsed redundant and closely related GO terms using the simplify() function with a similarity cutoff of 0.4.

### CytoTORCH Module Score Calculation

We passed a list of TORCH and cytokine receptors (Itgav, Dag1, Axl, Tyro3, Itgb1, Itgb3, Pdgfra, Pvrl1, Pvrl2, Mog, Tlr1, Tlr2, Ccr5, Tlr3, Tlr7, Tlr12, Cdh1, Met, Nlrp3, Cx3cr1, Cxcr4, Cxcr3, Cxcr2, Ccr1, Il17ra, Il1r1, Il1r2, Ifngr1, Ifngr2, Il4ra, Il6ra, Il15ra, Il2rb, and Il2rg) to Seurat’s AddModuleScore function which calculates the average expression levels for the requested genes subtracted by the aggregate expression of randomly selected control features (see Seurat documentation for details).

### Inferring transitions between substates over time in developing microglia

We used the FLOWMAPR package^54^ to infer cell state transitions over time in developing microglia. We used the top 12 principal components from our merged developing microglia dataset (described above) as input, and clustered the cells within each timepoint (E12.5-P1) into 693 nodes using FLOWMAPR’s k-means algorithm. We used FLOWMAPR in “single” mode to construct a force-directed graph based on Euclidean distance between these nodes, only allowing edges between nodes belonging to the same or consecutive timepoints. To determine whether transitions between each pair of substates were more likely than random chance, for each microglial substate we examined all possible edges originating from nodes with within that substate and terminating in valid nodes within other substates as a population for a series of Fisher’s exact tests: for each target substate, we defined categories for contingency tables based on whether each possible edge within the population terminates within the target substate, and whether each possible edge was actually found by FLOWMAPR. We visualized the odds ratios and p-values for transitions between each pair of microglial substates using the BioCircos package in R.

### Statistics and Reproducibility

To test whether MIA altered the relative proportions of each microglial substate, we used the *lme4* R package to construct binomial generalized linear mixed effect models. For each substate, we compared two models, each with a binary vector indicating whether each cell in the dataset belonged to that substate as the response variable and the litter each cell originated from as a random effect; one model included MIA treatment status as an additional fixed effect, the other did not. We used the anova function in R to calculate a p-value for the significance of including treatment as a fixed effect. We used identical methods to test for changes in cell type proportions between MIA and NaCl in WT and CSF1R-/- mice.

While no statistical methods were used to predetermine sample size, the sample sizes are similar to those used in the field^12,22,43^. Animals were randomized into treatment groups with similar numbers per group when possible. Investigators were not blinded during analysis of sequencing data due to the nature of the experiments. Statistical analyses were performed using R, and we controlled for multiple comparisons where appropriate using a corrected P value cutoff of <0.05. Information regarding specific statistical comparisons and packages is listed above.

## SUPPLEMENTARY MATERIALS

Fig. S1. Response to maternal immune activation

Fig. S2. Microglia isolation by FACS

Fig. S3. Ms4a7+ microglia characterization

Fig. S4. Proliferating microglia characterization.

Fig. S5. Embryonic microglia clusters after MIA.

Fig. S6. Gene ontology analysis in embryonic and postnatal microglia after MIA.

---

**Fig. S1.**
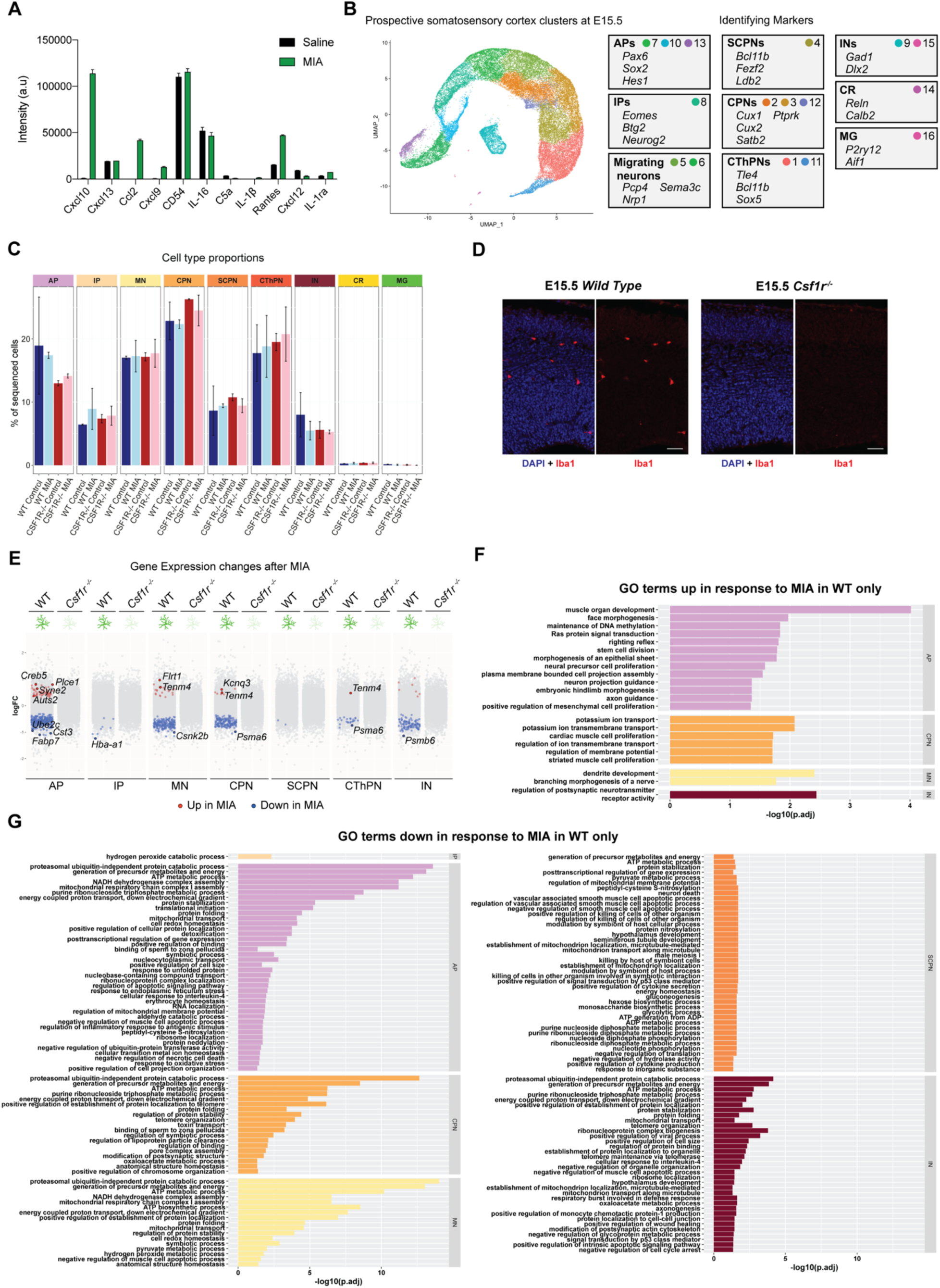
Response to maternal immune activation. (A) Bar plot showing cytokine expression analysis in spleens of pregnant dams 5h after being injected with saline or Poly(I:C) (MIA), measured via cytokine array profiling (see Methods). (B) UMAP visualization of snRNAseq from the developing somatosensory cortex at E15.5. All samples from WT and CSF1R KO, MIA and saline control-treated, are included. Cells are colored by cluster (left). Canonical marker genes used for cell type identification are shown on the right, along with the cluster numbers that correspond to each cell type. (C) Bar plot showing cell type proportions after maternal injection of saline as compared to Poly(I:C) in WT and CSF1R KO mice. Generalized mixed effects binomial model showed no significant changes in substate proportions between treatment groups. Error bars represent 1 standard deviation. (D) Representative images of Iba1 immunostaining in *Wild Type* and *Csf1r^-/-^* prospective somatosensory cortex at E15.5 (scale bar 50μm). (E) Dot plot of log(fold change) in gene expression between MIA and saline for cell types with significant changes at E15.5. DEG analysis performed using DESeq2. AP, apical progenitors; IP, intermediate progenitors; MN, migrating neurons; CPN, callosal projection neurons; SCPN, subcerebral projection neurons; CThPN, corticothalamic projection neurons; IN, interneurons. (F) Enriched Gene Ontology (GO) terms from differentially expressed genes upregulated in response to MIA in WT but not in CSF1R KO mice. (G) Enriched GO terms from differentially expressed genes downregulated in response to MIA in WT but not CSF1R KO mice.

**Fig. S2.**
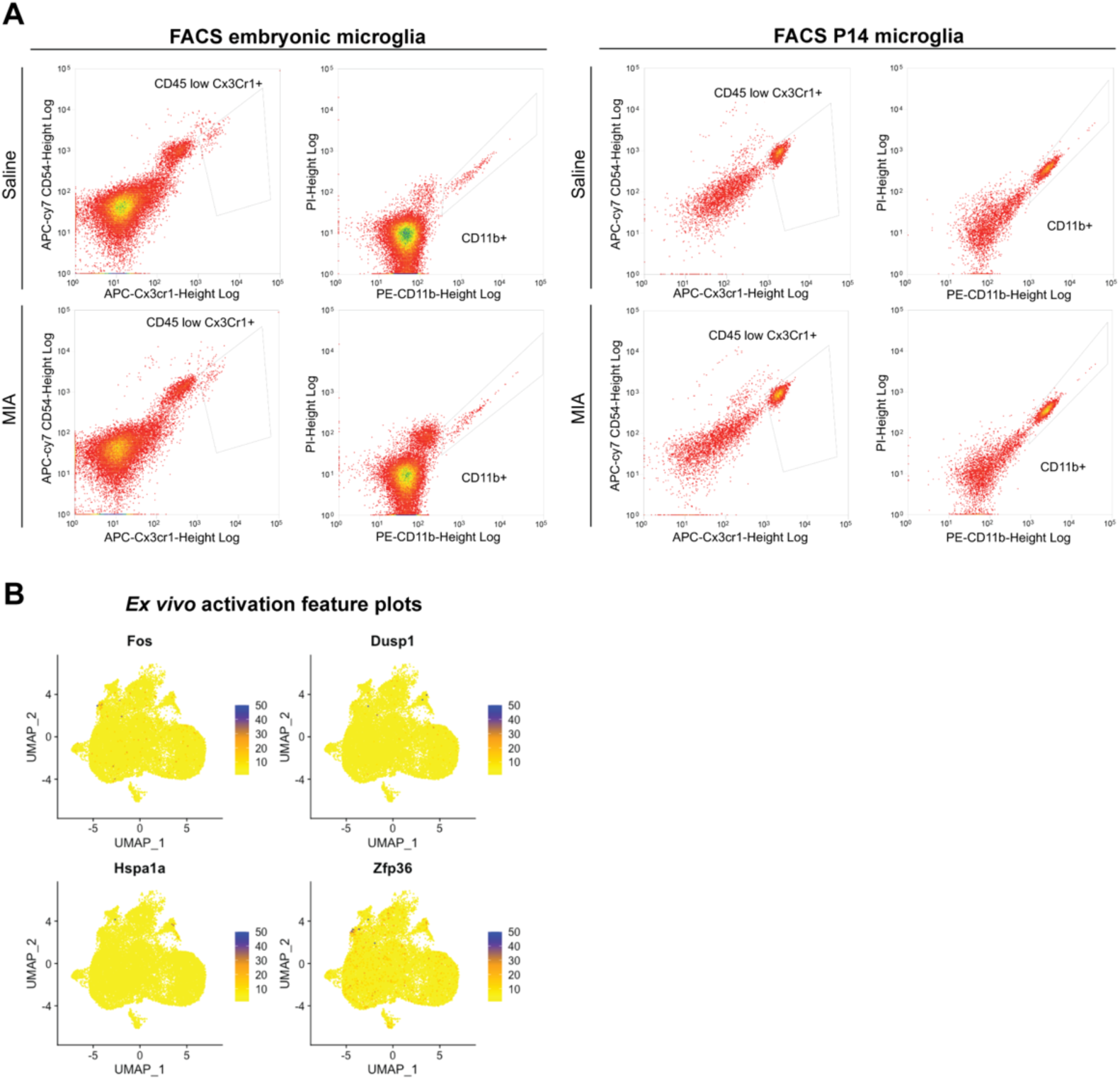
Microglia isolation by FACS. (A) FACS plots of microglia sorting in embryonic and postnatal day 14 (P14) mice after maternal injection of saline or Poly(I:C) (MIA). Prior to FACS purification, microglia were labeled with antibodies against CD45, CX3CR1 and CD11b. Boxes show gating for each parameter. (B) UMAP visualization of microglia scRNAseq data from the developing somatosensory cortex of WT mice. Plots are colored for expression of known microglia *ex vivo* activation markers.

**Fig. S3.**
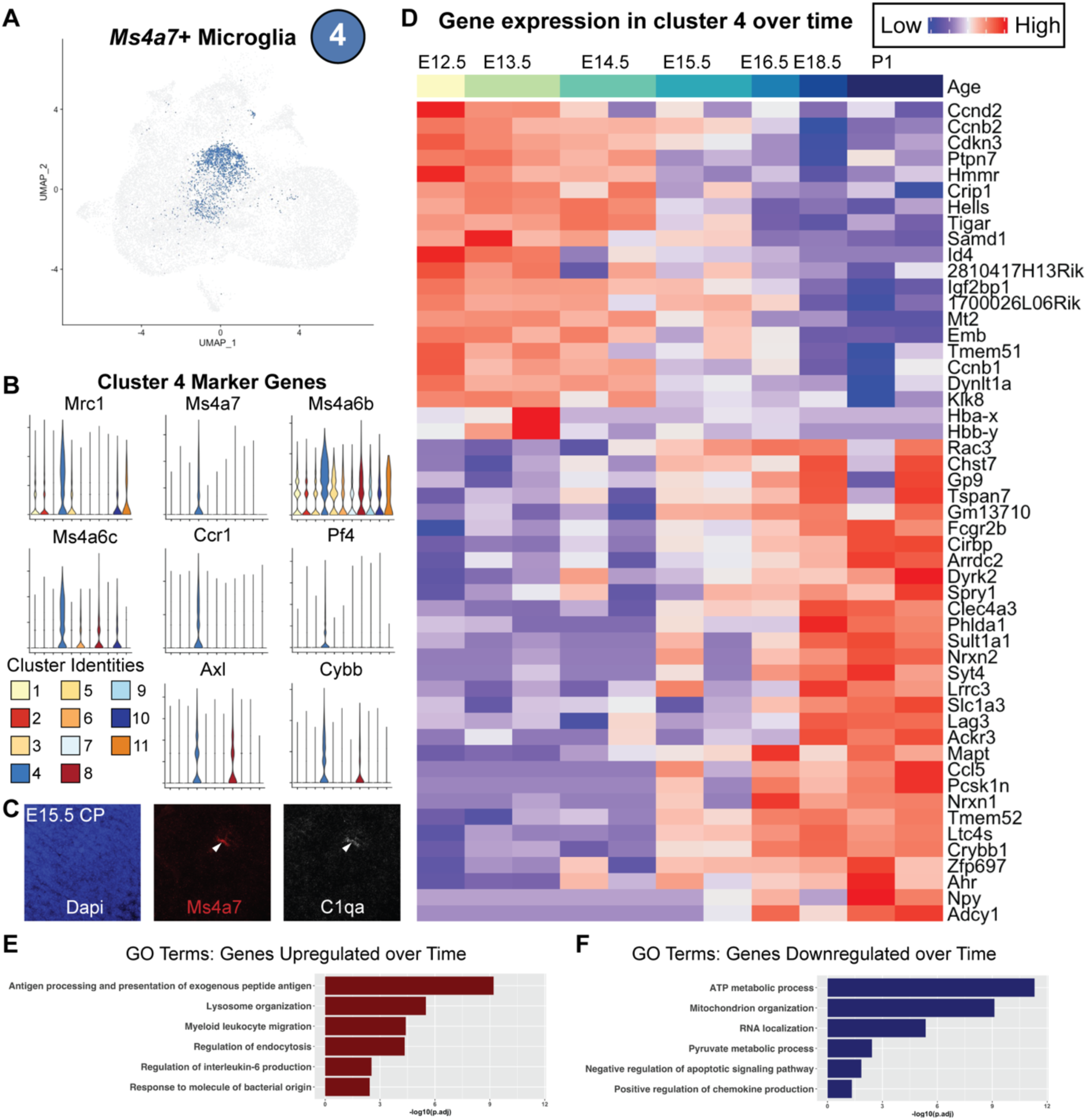
Ms4a7+ microglia characterization. (A) UMAP visualization of microglia scRNAseq data from the developing somatosensory cortex of WT mice. Cells belonging to microglia cluster 4 are colored. (B) Violin plots showing the expression of microglia cluster 4 marker genes in all clusters. (C) Fluorescent images from RNA *in situ hybridization* (RNAScope) showing colocalization of Ms4a7 and C1qa (pan-microglial marker) in the mouse cortical plate (CP) at E15.5. (D) Heat map of gene expression in cluster 4 over time. (E) Bar plots showing enriched Gene Ontology (GO) terms in genes upregulated over time in microglia cluster 4. (F) Bar plots showing enriched GO terms in genes downregulated over time in microglia cluster 4.

**Fig. S4.**
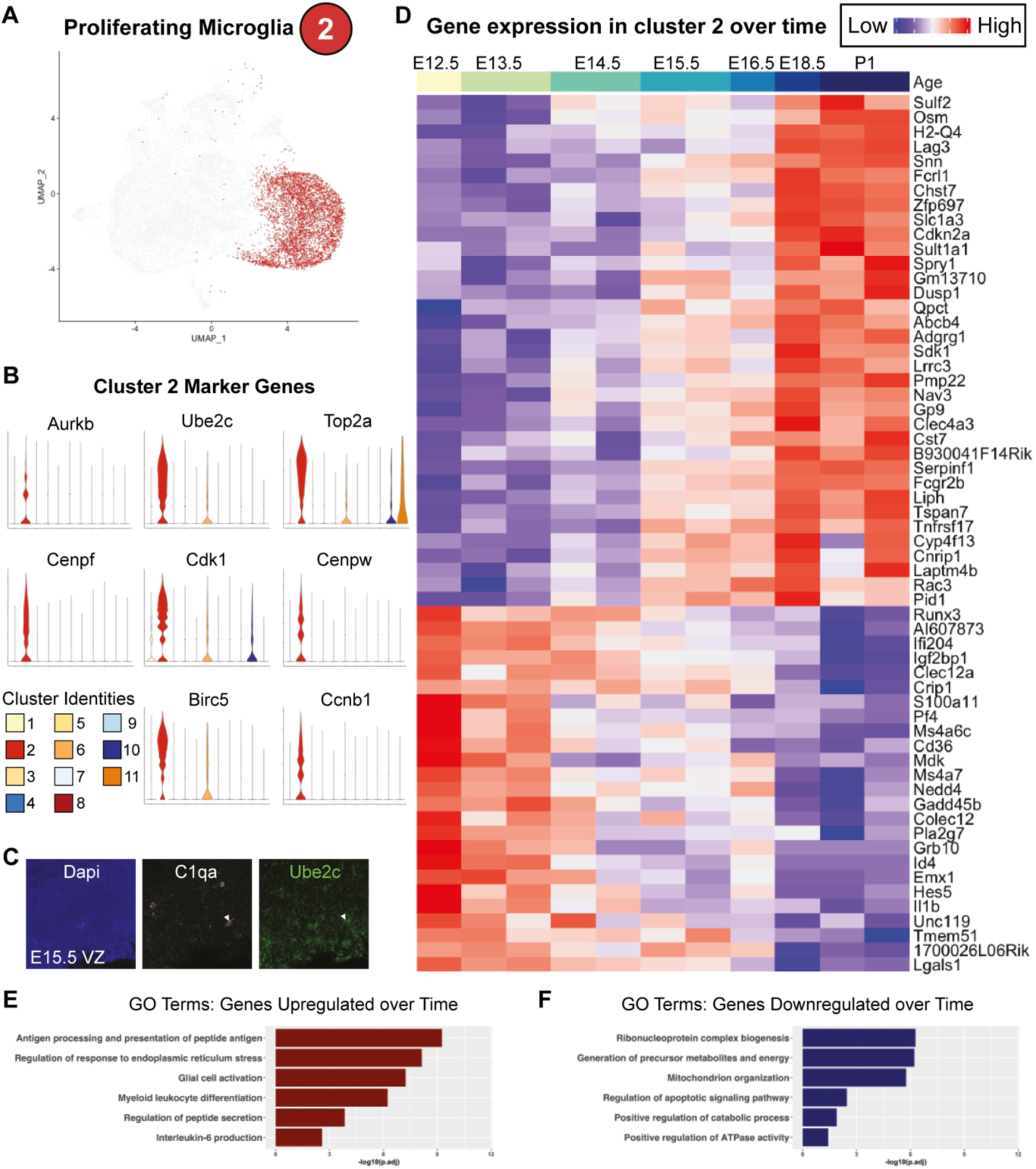
Proliferating microglia characterization. (A) UMAP visualization of microglia scRNAseq data from the developing somatosensory cortex of WT mice. Cells belonging to microglia cluster 2 are colored. (B) Violin plots showing the expression of microglia cluster 2 marker genes in all clusters. (C) Fluorescent images from RNA *in situ hybridization* showing colocalization of Ube2c and C1qa (pan-microglial marker) in the mouse ventricular zone (VZ) at E15.5. (D) Heat map of gene expression in cluster 2 over time. (E) Bar plots showing enriched Gene Ontology (GO) terms in genes upregulated over time in microglia cluster 4. (F) Bar plots showing enriched GO terms in genes downregulated over time in microglia cluster 4.

**Fig. S5.**
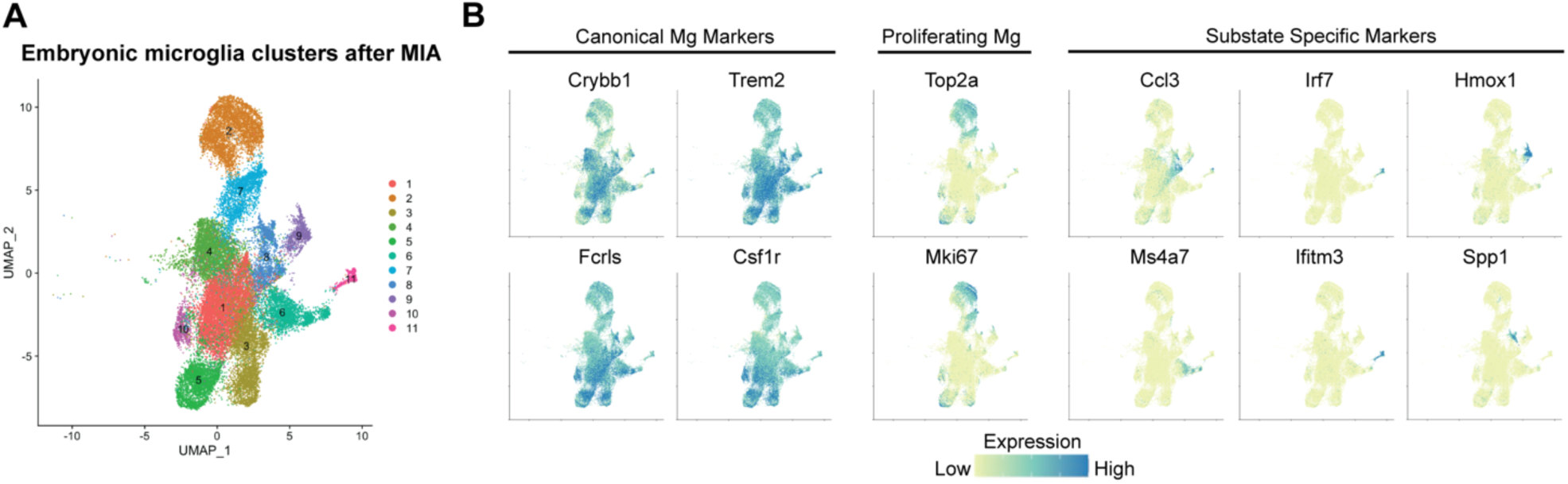
Embryonic microglia clusters after MIA. (A) UMAP visualization of microglia scRNAseq data from the developing somatosensory cortex of mice after maternal injection of Poly(I:C) (MIA). UMAP shows all cells from MIA and saline-injected animals analyzed at both E13.5 and E15.5. Cells are colored by cluster. (B) UMAP plots from A colored for expression of canonical, proliferating and substate specific microglia marker genes.

**Fig. S6.**
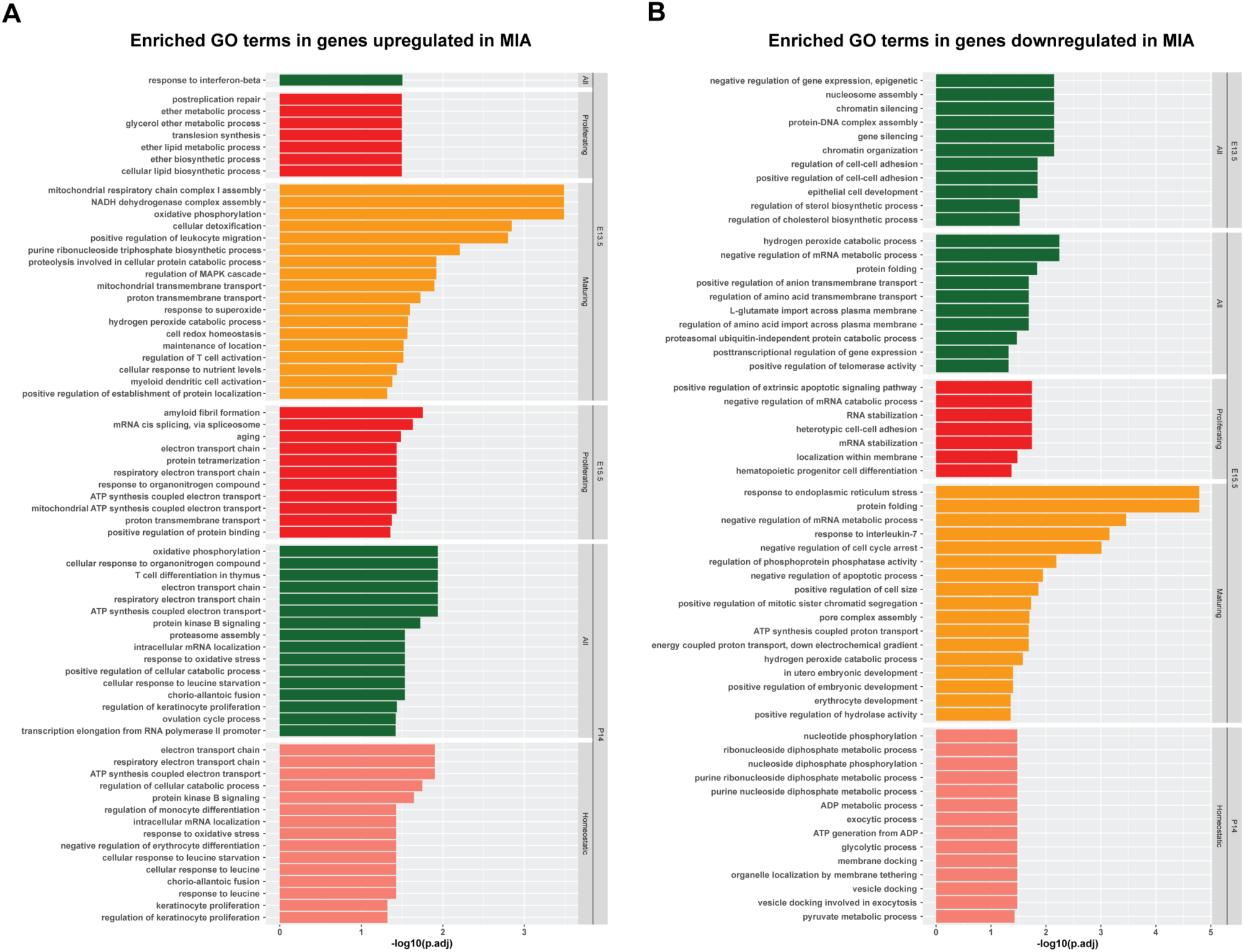
Gene ontology analysis in embryonic and postnatal microglia after MIA. (A) Bar plots showing enriched Gene Ontology (GO) terms from differentially expressed genes upregulated in response to MIA in WT mice. (B) Bar plots showing enriched GO terms from differentially expressed genes downregulated in response to MIA in WT mice.

